# Enhanced reproductive thermotolerance is associated with increased accumulation of flavonols in pollen of the tomato *high-pigment* 2 mutant

**DOI:** 10.1101/2020.12.18.423528

**Authors:** Nicholas Rutley, Golan Miller, Fengde Wang, Jeffrey F Harper, Gad Miller, Michal Lieberman-Lazarovich

**Author notes:** Institute of Vegetables and Flowers, Shandong Academy of Agricultural Sciences, Jinan, China (Fengde Wang).

## Abstract

Climate change has created an environment where heat stress conditions are becoming more frequent as temperatures continue to rise in crop production areas around the world. This situation leads to decreased crop production due to plant sensitivity to heat stress. Reproductive success is critically dependent on plants’ ability to produce functional pollen grains, which are the most thermo-sensitive tissue. Flavonols are plant secondary metabolites known for their potent antioxidative activity, essential for male fertility in several species including tomato, and implicated in heat stress tolerance. Since flavonols are highly abundant in fruits of the tomato *high pigment-2* (*hp2*) mutant, we tested the level of flavonols in pollen of this mutant, under the hypothesis that increased accumulation of flavonols would render pollen more tolerant to heat stress. Indeed, pollen from three alleles of the *hp2* mutant were found to have flavonols levels increased by 40% on average compared with wild-type under moderate chronic heat stress conditions. This mutant produced on average 7.8-fold higher levels of viable pollen and displayed better germination competence under heat stress conditions. The percentage of fully seeded fruits and the number of seeds per fruit were maintained in the mutant under heat stress conditions while decreased in wild-type plants. Our results strongly suggest that increased pollen flavonols content enhances pollen thermotolerance and reproductive success under heat stress conditions. Thus, the high flavonols trait may help frame the model for improving crop resilience to heat stress.

## Introduction

Crop productivity is often dependent on efficient sexual reproduction, a complex process that is very sensitive to environmental stresses, in particular heat stress, leading to considerable yield losses. Sensitivity to heat stress during the reproductive phase of plant development was documented in various crop species, and it was shown to be manifested by morphological alterations of anthers, style elongation, bud abscission and reduced fruit number, weight and seed set. The most heat stress sensitive processes include meiosis, pollen germination, pollen tube growth, pollen/pistil interactions, fertilization, formation of the endosperm and embryo development (Abdul-Baki and Stommel 1995; Firon et al. 2006; Peet, Sato, and Gardner 1998; Rudich, Zamski, and Regev 1977; Warrag and Hall 1984; Monterroso and Wien 1990; Erickson and Markhart 2002). Severity of the damage is dependent on the nature of the heat stress experienced by the plants (i.e., intensity, duration, occurrence of heat spikes etc.), and whether or not thermotolerance was acquired. In tomato (*Solanum lycopersicum*), a prolonged stress of day temperatures exceeding 32°C with night temperature above 20°C caused reduced fruit set, fruit weight, total yield and seed production (El Ahmadi and Stevens 1979; Peet et al. 1998). Prolonged exposure to temperature of 36°C led to the development of abnormal anthers and pollen grains, and style elongation (Rudich et al. 1977; Giorno et al. 2013). Predictions of the effect of temperature increment on major crops yield show that each degree-Celsius increase in global mean temperature would, on average, reduce yields by 3.1-7.4% (Zhao et al. 2017). Therefore, a comprehensive understanding of how plants respond to heat stress conditions to mitigate possible damage is fundamental for designing measures to sustain adequate productivity under increasingly warming climate.

### Pollen sensitivity to heat stress

Pollen development is considered the most heat-sensitive stage during plant development (Lohani et al. 2020), as it was shown to be more sensitive than both the sporophyte and female gametophyte (Peet et al. 1998; Young et al. 2004; Wang et al. 2019). Pollen viability was reported to be compromised by high temperatures in crop plants such as wheat (Begcy et al. 2018), rice (Jagadish et al. 2007), sorghum (Djanaguiraman et al. 2018), soybean (Djanaguiraman et al. 2013), and tomato (Firon et al. 2006). The occurrence of heat stress during male reproductive development, eight to thirteen days before anthesis, impairs pollen viability and function, leading to decreased fruit and seed set (Iwahori 1966; Peet et al. 1998; Zinn et al. 2010; De Storme and Geelen 2014; Muller and Rieu 2016). Heat stress causes disruption of meiotic cell division, abnormal pollen morphology and size, and reduced grain number, viability, and germination capacity (Endo et al. 2009; Peet et al. 1998; Djanaguiraman et al. 2013; Giorno et al. 2013; Pressman et al. 2002; Firon et al. 2006; Begcy et al. 2019; Prasad et al. 2006). In tomato, optimal growth conditions are approximately 26-28°C during the day and 20-22°C during the night. Pollen heat stress related damage was observed after short episodes of high temperatures at 40°C, as well as after chronic exposure to milder heat stress of 32°C/28°C or 31°C /25°C day/night for several months (Firon et al. 2006; Iwahori 1966).

### Flavonols in pollen development and heat stress tolerance

Flavonols are plant secondary metabolites, among the most abundant groups of flavonoids, which are present in large quantities in various tissues of many plant species (Buer et al. 2010). Flavonols are important for stress defense and proper pollen function (Falcone Ferreyra et al. 2012), being essential for maintenance of pollen fertility. Mutants deficient in chalcone synthase (CHS), the first dedicated enzyme in the flavonol biosynthesis branch, cannot produce pollen tubes. The infertility phenotype of this mutant can be rescued by exogenous application of flavonols like quercetin or kaempferol (Mo et al. 1992; Ylstra et al. 1992; Falcone Ferreyra et al. 2012).

Pollen flavonols are synthesized in the tapetum, the innermost cell layer of the anther, and transported to microspores during pollen development, becoming a major component of the pollen coat (Hsieh and Huang 2007; Fellenberg and Vogt 2015). It has been suggested that flavonols might also contribute to pollen wall plasticity to facilitate rapid pollen tube growth (Derksen et al. 1999).

Flavonols are potent antioxidants that have been implicated in abiotic stress tolerance because they can inhibit the generation of free radicals and scavenge reactive oxygen species (ROS) (Brown et al. 1998; Rice-Evans 2001; Melidou et al. 2005; Agati et al. 2012; Brunetti et al. 2013; Martinez et al. 2016). In particular, several studies pointed at the involvement of flavonols in the response to heat stress. In tomato, flavonols content in leaves and pollen was increased in response to heat stress (Martinez et al. 2016; Paupière et al. 2017). In leaves, heat stress specific activation of the flavonol-biosynthesis pathway was observed, suggesting a specific role for flavonols in mitigating the oxidative stress developed under high temperatures (Martinez et al. 2016). Using a complementary approach, a non-targeted metabolomics study in pollen suggested that the most significant metabolic response to a two hours heat stress was the two-fold increase in total flavonoids, which include flavonols (Paupière et al. 2017). In addition, several studies correlated the reduction of flavonols with a reduction in heat stress tolerance. For example, a CHS-deficient mutant variety of Morning Glory (*Ipomoea purpurea*), which does not produce flavonols, shows reduced pollen fertility under high temperatures (Coberly and Rausher 2003). In tomato, knockdown of Heat shock factor A2 (HsfA2) resulted in reduced pollen viability in response to heat stress and decreased chalcone isomerase (CHI) expression (Fragkostefanakis et al. 2016). The tomato anthocyanin reduced (*are*) mutant has reduced flavonol levels in pollen grains and tubes. This mutant produce fewer pollen grains and is impaired in pollen viability, germination, tube growth, and tube integrity, resulting in reduced seed set. Consistent with flavonols acting as ROS scavengers, *are* shows elevated levels of ROS in pollen tubes and inhibited tube growth that could be rescued by antioxidant treatment (Muhlemann et al. 2018). That study showed that flavonols support pollen development and tube growth under heat stress by reducing the abundance of ROS.

### The tomato *high pigment 2* mutant

The pleiotropic *high-pigment 2* (*hp2*) natural mutant in tomato is characterized by enhanced light responsiveness resulting in increased pigmentation and increased plastid number and size (Mustilli et al. 1999; Levin et al. 2003; Kolotilin et al. 2007). Metabolomic studies revealed that the darker red color of ripe fruits of the *hp2* mutant is caused by increased levels of secondary metabolites, mainly carotenoids and flavonoids. Total carotenoids in *hp2* fruits were increased up to 3.2-fold (Kolotilin et al. 2007) and the flavonoids rutin and a quercetin derivative were found to be induced 2.9 and 4.4-fold, respectively (Bino et al. 2005).

The gene underlying *hp2* is the tomato homologue of the Arabidopsis *DEETIOLATED1* (*DET1*) gene (Solyc01g056340), initially identified as a negative regulator of photomorphogenesis (Levin et al. 2003; Mustilli et al. 1999; Pepper et al. 1994). Three *hp2* alleles are known: *hp2, hp2*^*j*^ and *hp2*^*dg*^. In *hp2*, a nine base pairs deletion within a putative nucleolar localization signal of *DET1* results in a three amino acids deletion. *hp2*^*j*^ and *hp2*^*dg*^ are the result of a conserved amino acid substitution within a protein-protein interaction domain (Mustilli et al. 1999; Jones et al. 2012). These alleles share similar phenotypes but also differences, as for example, the *hp2*^*dg*^ allele has more intense pigmentation at the green fruit stage than *hp2* (Konsler 1973; van Tuinen et al. 1997) while *hp2*^*j*^ has shorter hypocotyls and higher hypocotyl anthocyanin content than *hp2* (Mustilli et al. 1999; Levin et al. 2003). While *DET1* is expressed throughout the tomato fruit in almost all tissues and ripening stages tested (http://tea.solgenomics.net), no major transcriptional changes were reported for *DET1* in the *hp2* mutants. Downregulation of *DET1* specifically in tomato fruit resulted in a significant increase in carotenoids, flavonoids, and chlorophyll levels, along with increase in plastids number and area per cell (Davuluri et al. 2005; Enfissi et al. 2010), supporting earlier studies that define DET1 as regulator of secondary metabolites biosynthesis.

DET1 is a component of an E3 ligase complex involved in protein ubiquitination and degradation by the 26S proteasome (Bernhardt et al. 2006; Zhang et al. 2008). Among the proteins shown to be targets of the DET1 complex are positive regulators of photomorphogenesis such as *ELONGATED HYPOCOTYL5* (*HY5*) in Arabidopsis (Osterlund et al. 2000) and upstream regulators of chloroplast biogenesis, flavonoid accumulation and carotenoid biosynthesis such as *GOLDEN2-LIKE* (GLK2), BBX20 and MBD5 in tomato (Tang et al. 2016; Powell et al. 2012; Nguyen et al. 2014; Xiong et al. 2019; Li et al. 2016). Interestingly, ubiquitin ligases which contain DET1 as a complex unit, were suggested to play a role in abiotic stress responses as they mediate ABA signaling in Arabidopsis (Irigoyen et al. 2014; Seo et al. 2014). The involvement of DET1 in abiotic stress tolerance was further demonstrated by improved seeds germination of the *det1* Arabidopsis mutant under salt and osmotic stress conditions (Fernando and Schroeder 2016). However, the possible effect of the tomato *det1* (*hp2*) mutant on abiotic stress response or stress tolerance was not evaluated prior to this study. Taking into account that the *hp2* mutant has increased content of antioxidants capable of counteracting ROS produced under stress, we hypothesized that *hp2* may be more tolerant to heat stress. Considering the high sensitivity of pollen to heat stress, such hypothesis requires that pollen of *hp2* would also be enriched with antioxidants, not only leaves and fruits. This information was not available prior to our study. We therefore set to investigate pollen characteristics of *hp2*, i.e. pollen flavonols content and pollen viability, under conditions of heat stress, a major abiotic stress in tomato.

In this study, we compared the reproductive success of *hp2* tomato mutants and wild-type isogenic lines under moderate chronic heat stress (MCHS) conditions (34/24°C day/night temperatures for 10-12 weeks during the plants’ reproductive stage), and measured pollen viability and flavonols level at the population scale using flow cytometry. We show that the *hp2* mutants are more heat stress tolerant than their isogenic wild-type lines producing more seeds than the wild-types under MCHS, correlated with better pollen viability and higher pollen flavonols level. We also show that tomato cultivars known for being thermotolerant have increased flavonols content in pollen. Our findings indicate that the ability of pollen to accumulate high flavonols content can be employed as a marker predicting increased pollen thermotolerance and improved reproductive success under heat stress conditions. Moreover, our results may be translated into a biotechnology approach to enhance pollen flavonols levels in order to achieve heat stress tolerance in tomato and other crops.

## Results

### The *high pigment 2* mutant maintains normal seeds production under moderate chronic heat stress conditions

One of the most prominent effects of heat stress on reproductive development is reduced seed set (Abdul-Baki 1991; Sato et al. 2000; Xu et al. 2017). To test whether the *hp2* mutant performs better than the wild-type under heat stress conditions, the three allelic lines, *hp2, hp2*^*j*^ and *hp2*^*dg*^, were grown in temperature-controlled greenhouses along with their corresponding isogenic wild-type lines; Moneymaker (for *hp2, hp2*^*j*^) and Manapal (for *hp2*^*dg*^). The plants grew under control temperatures of 26/20°C day/night in two greenhouses until the onset of flowering. From that stage throughout the rest of the experiment, the temperature in one of the greenhouses was set to reach at least 32°C during the day and at least 22°C during the night, creating conditions of Moderate Chronic Heat Stress (MCHS), while the other greenhouse was maintained at control conditions (Figure S1a). The impact of MCHS was evident, as flower damage such as elongated style, abnormal anthers and flower drop was prevalent in the stress greenhouse compared with the control greenhouse (Figure S1b). Heat stress perception was confirmed at the molecular level by quantifying the expression level of *Hsp17*.*6* gene (NM_001246984.3) as a heat stress reporter (Frank et al. 2009). Under MCHS conditions, *Hsp17*.*6* was induced between 31 to 43-fold, depending on the genotype, relative to control conditions (Figure S1c).

To evaluate reproductive performance of the *hp2* mutant we estimated the rate of fully seeded fruits (i.e., having a full seed set by visual inspection, Figure 1a), and counted the number of seeds per fruit under both MCHS and control conditions (Figure 1c). Both wild-type lines, Moneymaker and Manapal, had a significant reduction of 5.4-fold and 4-fold in the proportion of seeded fruits per plant under MCHS conditions, respectively, whereas in all three *hp2* genotypes the proportion of seeded fruit was similar between control and MCHS conditions (Figure 1b). The number of seeds per fruit was very similar among all genotypes under control conditions, ranging from 56 to 80 seeds per fruit. Under MCHS conditions, the *hp2* genotypes maintained a similar number of seeds (79-87 seeds per fruit) whereas a decrease to 44 and 52 seeds per fruit was observed in the Moneymaker and Manapal genotypes, respectively (Figure 1c). The apparent reduction in fruit size of *hp2* under MCHS conditions compared with control conditions (Figure 1a) was addressed by measuring single fruit weight, but the effects were found to be non-significant (Figure 1d).

**Figure 1.**
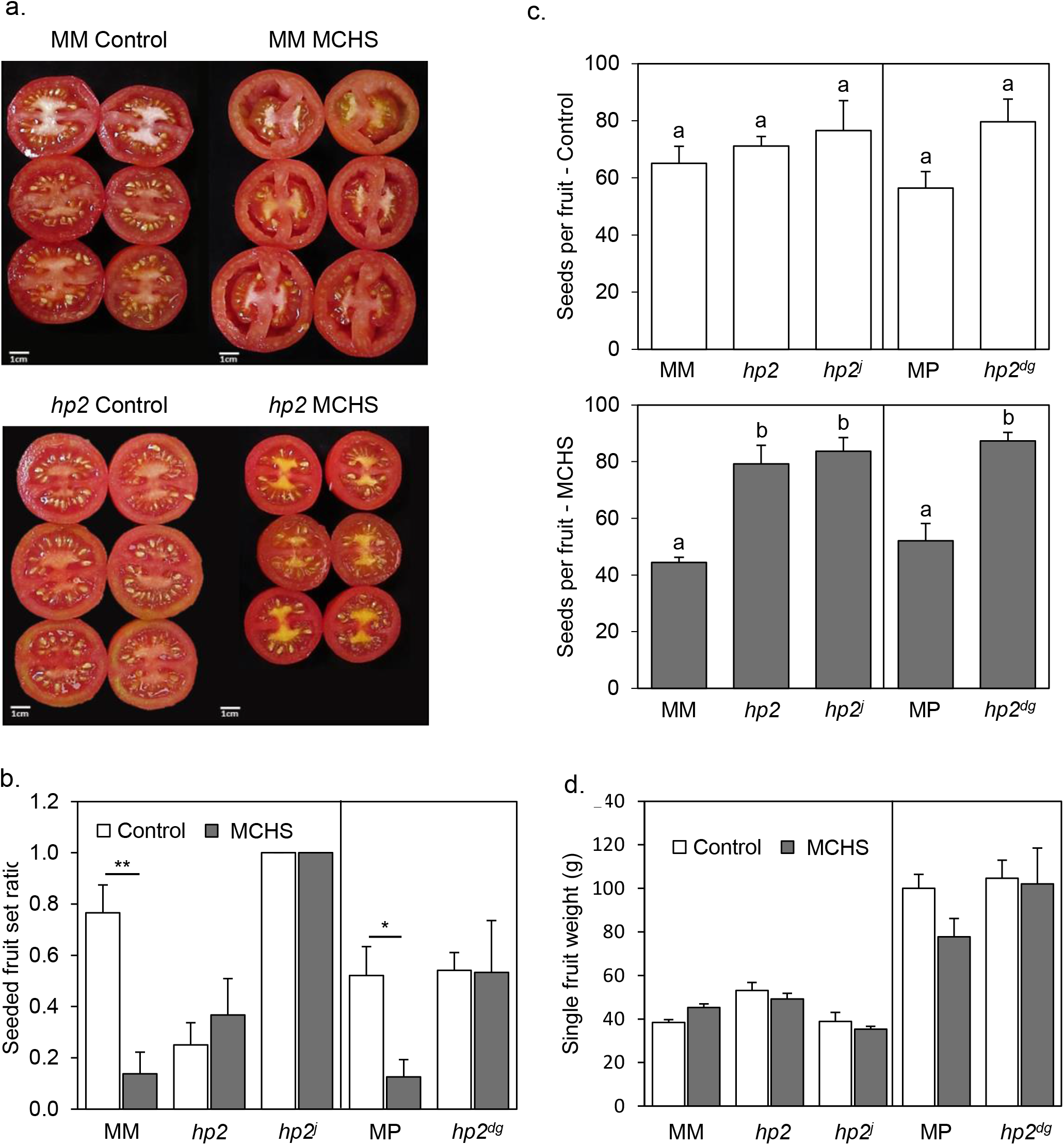
The *hp2* mutant maintains normal seed production under MCHS conditions. (a) Visual differences in fruit seed set and hollowness between *hp2* and moneymaker under control and MCHS conditions. (b) Rates of fruits full of seeds, under control and MCHS conditions. Fruits were scored as ‘full of seeds’ when visually similar to fruits shown in (a) for control conditions. (c) Number of seeds per fruit under control (upper panel) and MCHS conditions (bottom panel). (d) Single fruit weight under control and MCHS conditions. *, p-value < 0.05. **, p-value < 0.01. Different letters indicate statistically significant differences. MCHS, moderate chronic heat stress. MM, Moneymaker. MP, Manapal.

### The *high pigment 2* mutant maintains higher pollen viability and metabolic activity under moderate chronic heat stress conditions

Since the seed set of all three *hp2* mutants was not affected by heat stress conditions, we assumed that the *hp2* pollen might be more tolerant than the wild-type pollen. Therefore, we evaluated pollen viability under normal and MCHS conditions in *hp2* and wild-type lines using the 2’,7’-dichlorodihydrofluorescein diacetate (H_2_DCFDA) staining coupled with flow cytometry approach, as recently described (Luria et al. 2019; Rutley and Miller 2020). DCF (dichlorofluorescein) is the fluorescent molecule produced by ROS-mediated oxidation of H_2_DCFDA in viable cells and is an indication of ROS levels and metabolic activity (see more details in the experimental procedures section). Using this method, three pollen populations were detected: dead pollen, which do not produce a DCF signal (DCF-negative), and two populations of viable pollen, one having low metabolic activity (‘low ROS’) and the other with high metabolic activity (‘high ROS’) (Figure 2a). Flow cytometry analysis revealed similar distribution of the three pollen populations between the wild-type and *hp2* lines under control conditions with 71.1-79.9% viable pollen (low and high-ROS) across all genotypes. Under MCHS conditions, the percentage of viable pollen was dramatically reduced in all genotypes. However, the reduction was larger in the wild-type lines (0-2% high-ROS pollen fraction) compared with the *hp2* lines (3-18% high-ROS pollen fraction). Similarly, the low-ROS viable pollen fraction was bigger in the *hp2* lines (11.1-15.3%) compared with the wild-types (1.6-2.8%) under MCHS conditions. Overall, the viable pollen fraction (low and high-ROS) reached 1.6-4.4% of total pollen for the wild-types and 13.8-30.3% in *hp2* (Figure 2b, c, S2).

**Figure 2.**
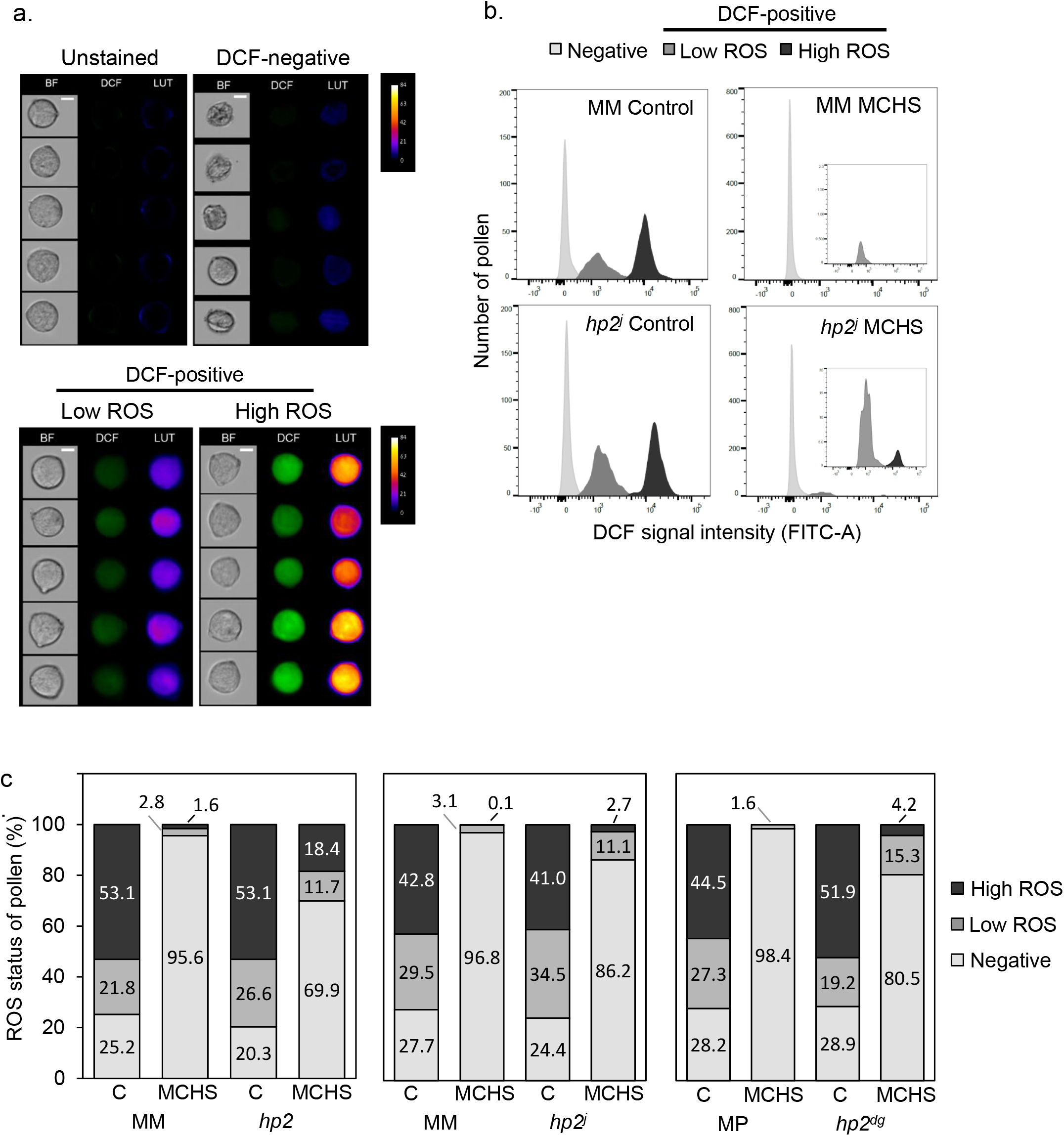
The *hp2* mutant maintains higher pollen viability and metabolic activity under MCHS conditions. (a) Randomly selected pollen grains from unstained and H_2_DCFDA-stained samples of Moneymaker wild type line under control conditions captured using the flow cytometer detecting DCF fluorescence. Pseudo-color lookup table (LUT) images were generated in ImageJ to emphasize the scale of DCF fluorescent intensity. BF, Brightfield images. Scale bar = 10 µm (b) Distribution of pollen by DCF-signal intensity. Inset boxes zoom into the DCF-positive fraction, which is smaller under MCHS conditions in all genotypes. (c) Distributions of ROS status (negative, low or high) of pollen across genotypes and conditions. Numbers denote the percentage of each status fraction. MCHS, moderate chronic heat stress. C, control conditions. MM, Moneymaker. MP, Manapal.

Temperature stress causes defects in developing pollen, often leading to deformed pollen and decreased grain size (Mercado et al. 1997; Porch and Jahn 2001). In flow cytometry, the forward scatter area (FSC-A) value is proportional to cell size (Adan et al. 2017). The distribution of pollen by FSC-A values revealed two pollen subpopulations, of smaller-grain and larger-grain size (Figure 3a, b). Under normal conditions, the fraction of larger pollen was bigger in either mutant or wild-type lines. On the contrary, under MCHS conditions, the majority of pollen grains were of smaller size, demonstrating heat stress damage. Additionally, under MCHS conditions, the larger-pollen faction was bigger in the *hp2* mutant compared with the related wild-type, implying better thermotolerance, in line with our results on pollen viability (Figure 3b, S4). Indeed, positive correlation was found between ROS level of viable pollen and grain size across all samples, meaning that high-ROS pollen were overall larger than low-ROS pollen (Figure 3c, Table S2). The high-ROS larger pollen fraction did not differ between the mutant and wild-type lines under normal conditions, however, under MCHS conditions, both *hp2* and *hp2*^*j*^ alleles produced 4 and 7-fold higher proportion of larger high-ROS pollen compared with the Moneymaker background. In the Manapal genotype, the larger high-ROS pollen was completely obliterated by the stress, whereas in the *hp2*^*dg*^ allele 14.8% of larger high-ROS pollen survived (Figure 3d).

**Figure 3.**
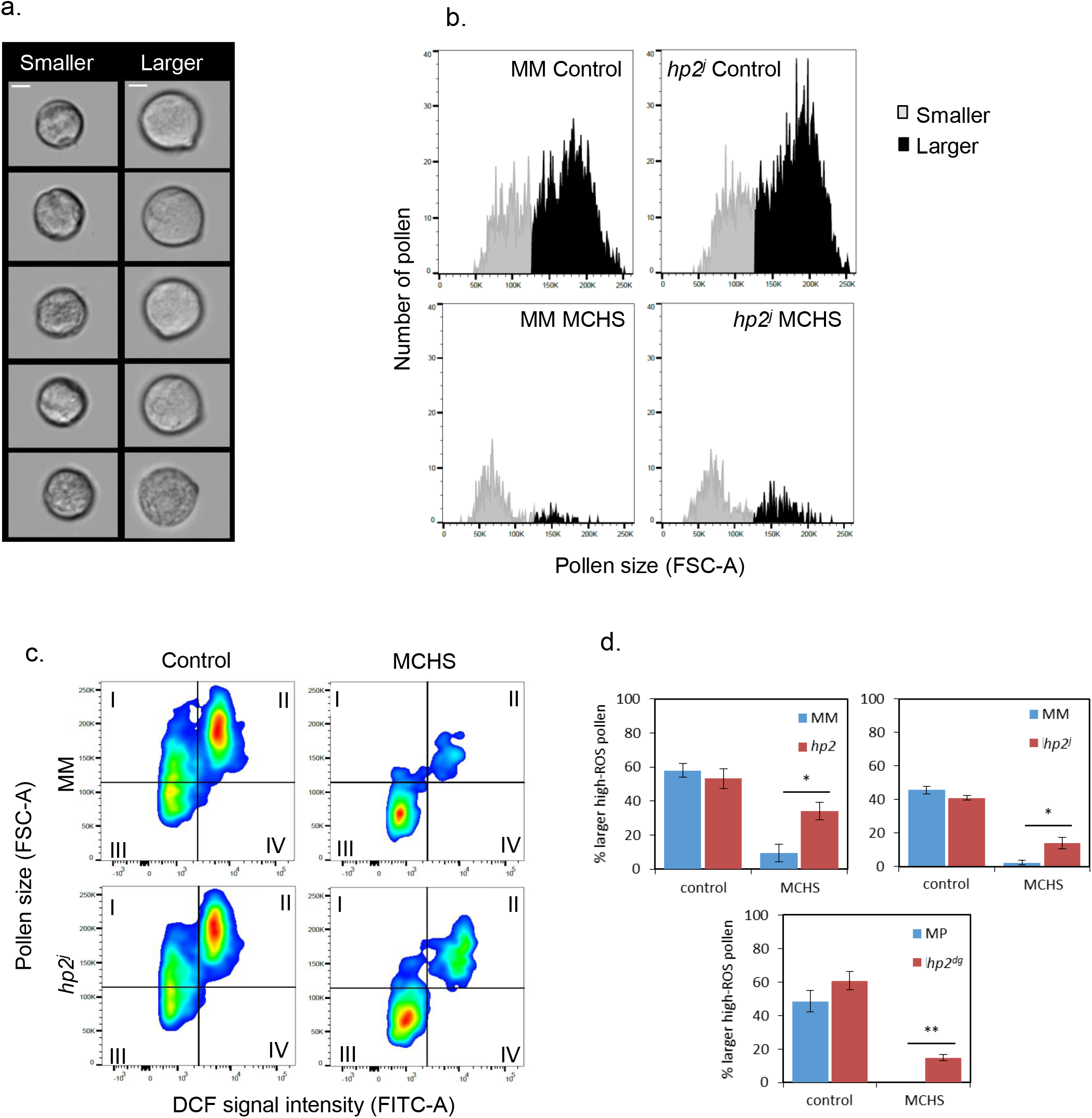
The *hp2* mutant maintains a higher proportion of larger pollen under MCHS conditions. (a) Randomly selected pollen grains from smaller and larger subpopulations in a Moneymaker control sample. Scale bar = 10 µm. (b) Pollen grain size distribution by FSC-A in an *hp2*^*j*^ sample from control and MCHS conditions. (c) DCF-positive pollen populations plotted by grain size (FSC-A, y-axis) versus DCF signal intensity (FITC-A, x-axis). The larger, high-ROS pollen faction is represented in quadrant II. (d) Rate of larger high-ROS pollen sub-population (section c - quadrant II) in all genotypes under control and MCHS conditions. *, p-value < 0.05. **, p-value < 0.01. MCHS, moderate chronic heat stress. C, control conditions. MM, Moneymaker. MP, Manapal.

### The *high pigment 2* mutant shows increased thermotolerance in terms of pollen germination

To ask whether the improved viability of *hp2* pollen under heat stress conditions could support thermotolerance during fertilization, we evaluated pollen germination *in vitro* under control and heat stress conditions. To this end, pollen was imbibed in germination medium and exposed to moderate heat stress conditions of 34°C for 30 minutes and the proportion of germinated pollen grains was scored following 1.5 hours of recovery at 22°C. We compared pollen of the MicroTom *hp2*^*dg*^ introgression line (MT *hp2*^*dg*^) (Sestari et al. 2014) with its MicroTom background line. Under control growth conditions (22°C) pollen germination rate was nearly two-fold higher in *hp2*^*dg*^ pollen compared with the wild-type MicroTom (MT) line. Heat stress led to a 12.5-fold and a 5.9-fold reduction in pollen tube production rate in the MT and the *hp2*^*dg*^ lines (corresponding to 2.5% and 9.9% germination), respectively (Figure 4a, b).

**Figure 4.**
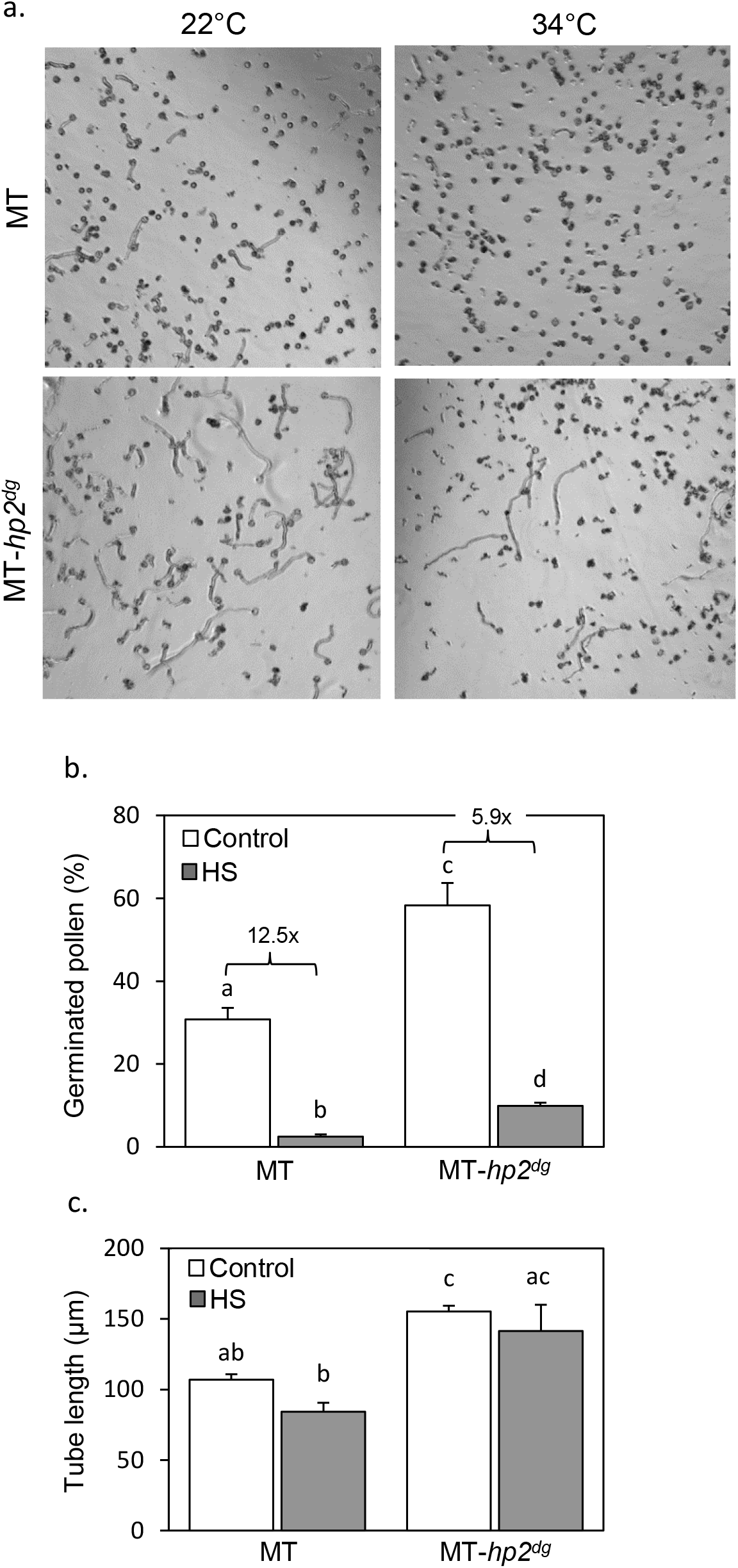
The *hp2* mutant shows better pollen germination following *in-vitro* heat stress treatment. (a) Germinated pollen of *hp2*^*dg*^ and MicroTom background following an *in-vitro* heat stress treatment of 34°C for 30 minutes, compared with germination without heat stress (22°C). (b) Percentage of germinated pollen for *hp2*^*dg*^ and MicroTom background following an *in-vitro* heat stress treatment, compared with control conditions. Numbers denote fold change between control and heat stress samples in each genotype. (c) Pollen tube length of *hp2*^*dg*^ and MicroTom pollen following an *in-vitro* heat stress treatment, compared with control conditions. Different letters indicate statistically significant differences. HS, heat stress. MT, MicroTom.

In addition, we measured the average length of pollen tubes and found that under both control and *in-vitro* heat stress, *hp2*^*dg*^ pollen had longer pollen tubes than wild-type indicating faster growth rate (Figure 4c). The heat stress treatment had a milder impact on tube length compared with pollen germination proportion, yet in the MT line tube length was reduced by 21% while in *hp2*^*dg*^ we measured 10% reduction (Figure 4c). Taken together, these results suggest that the increased thermotolerant in the *hp2*^*dg*^ line during tube production and growth contributed to the overall improved reproductive heat stress tolerance of the *hp2* lines *in vivo* (Figure 1).

### The *high pigment 2* mutant accumulates higher levels of flavonols in pollen developed under moderate chronic heat stress conditions

Considering the antioxidative property of flavonols and their established role in mitigating stress damage, the high level of flavonols in fruits of the *hp2* mutant (Bino et al. 2005) prompted us to test the levels of flavonols in *hp2* pollen. To this end, we stained mature pollen with the flavonol-specific stain diphenylboric acid-2-aminoethyl ester (DPBA) (Sheahan and Rechnitz 1992) and analyzed the fluorescence intensity using flow cytometry (Figure 5a). Signal intensity corresponds to levels of the major flavonols found in pollen, kaempferol and quercetin. Under control conditions, the mean fluorescence intensity (MFI) of the DPBA signal was marginally higher in the *hp2* lines compared with the wild-type lines. However, the DPBA signal significantly increased in all *hp2* lines under MCHS conditions, compared with only minor increase in the wild-type lines (Figure 5b). At the population level, we noticed that the distribution of pollen MFI values differed between wild-type and *hp2* lines and between control and MCHS conditions (Figure 5c). We quantified the fraction of pollen which hyper-accumulated flavonols (enhanced DPBA signal) and found that under MCHS conditions, the percentage of pollen which hyper-accumulated flavonols was higher in the *hp2* lines compared with their corresponding wild-type lines (Figure 5d). In *hp2*, the percentage of enhanced DPBA pollen was 5.8% on average, which translate to 1.75-fold difference compared with the Moneymaker wild-type under the same conditions (3.3%). *hp2*^*j*^ presented the strongest effect, reaching 35% of enhanced DPBA pollen under MCHS conditions, a 9.2-fold difference compared with the Moneymaker wild-type (3.8%). The weaker effect in *hp2*^*dg*^, a non-significant 1.4-fold higher in the mutant compared with the wild-type under MCHS conditions, may be attributed to the different genetic background compared with *hp2* and *hp2*^*j*^ (Figure 5d).

**Figure 5.**
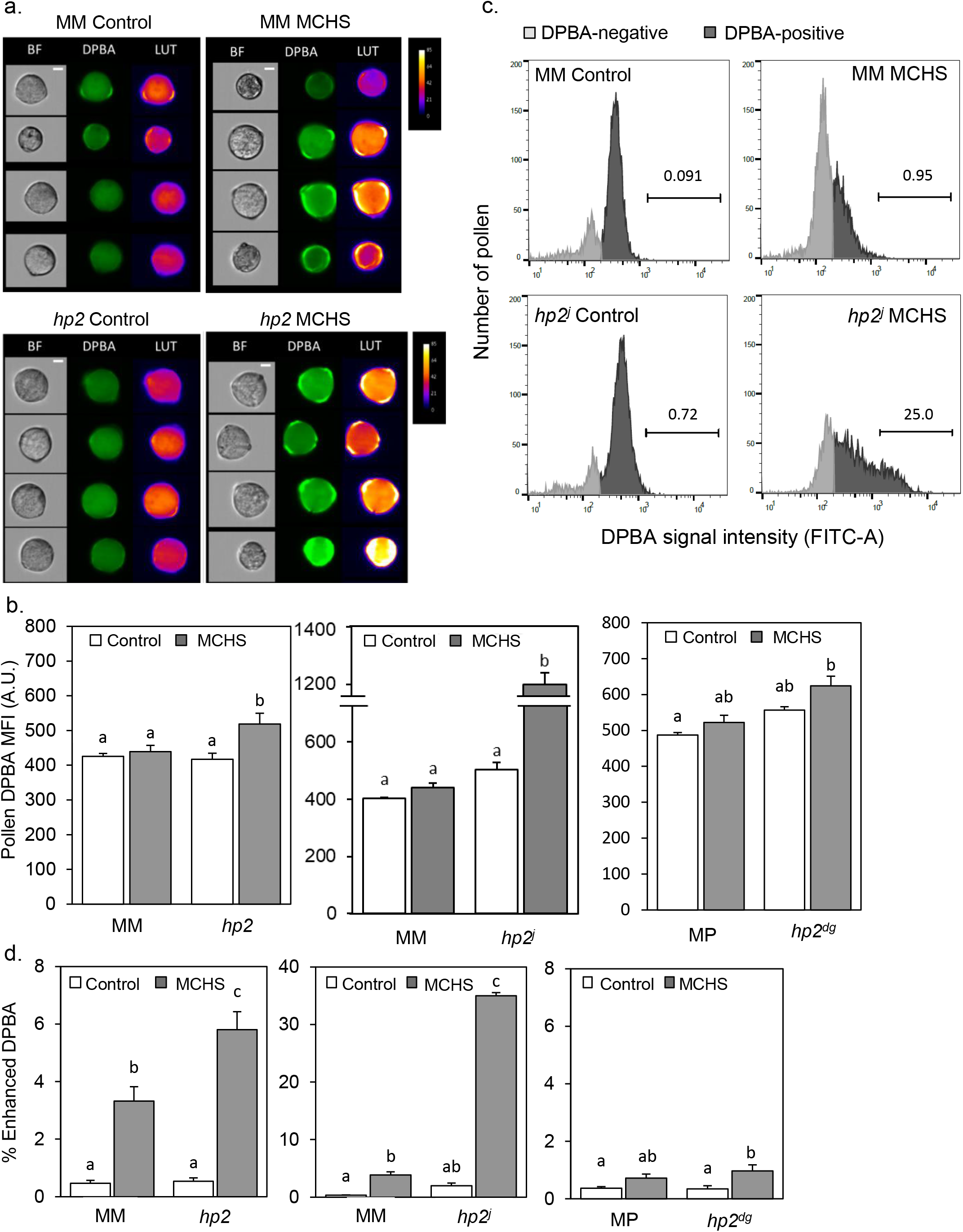
The *hp2* mutant accumulate higher levels of flavonols in pollen in response to MCHS. (a) Randomly selected pollen grains from DPBA-stained samples of *hp2* and Moneymaker under control and MCHS conditions. Pseudo-color lookup table (LUT) images indicate the scale of DCF fluorescent intensity. BF, Brightfield images. Scale bar = 10 µm (b) Pollen flavonol levels under control and MCHS conditions by DPBA stain mean fluorescence signal intensity (MFI) presented in arbitrary units. (c) Pollen distribution by DPBA signal intensity in *hp2*^*j*^ samples from control and MCHS conditions. Horizontal bars mark the enhanced-DPBA pollen subpopulation and the above number denotes the percentage of this subpopulation out of total DPBA-positive pollen. (d) Percentage of enhanced DPBA (i.e. enhanced flavonols) subpopulation under control and MCHS conditions. Different letters indicate statistically significant differences. MCHS, moderate chronic heat stress. MM, Moneymaker. MP, Manapal

Altogether, we show that *hp2* accumulates higher levels of flavonols in pollen, and that its content increases in response to MCHS conditions. We therefore hypothesize that flavonols accumulation contribute to the observed pollen thermotolerance of *hp2*. These results suggest that flavonols take part in the response to heat stress more profoundly in the *hp2* mutant.

### Flavonols levels are higher in pollen of heat stress tolerant tomato cultivars

To support the hypothesis that increased levels of flavonols is contributing to pollen thermotolerance, we compared the level of flavonols in pollen of two known heat stress tolerant cultivars, Saladette and CLN1621L, with two heat stress sensitive cultivars, M82 and Moneymaker.

The four cultivars (Saladette, CLN1621, M82 and Moneymaker) were grown in a non-controlled greenhouse during the summer hence reaching high-temperature conditions of 45/23°C day/night during flowering and fruit set. As a result, style elongation, a characteristic heat-stress phenotype, was abundant in flowers of M82 and Monemaker. In the heat stress tolerant cultivars CLN1621 and Saladette, most of the flowers appeared normal and undamaged (Figure 6a). The stress response was further validated by testing the expression of the heat stress molecular marker gene *Hsp17*.*6*. As expected, this gene was induced under heat stress relative to normal conditions, with the tolerant cultivars Saladette and CLN1621 showing no or non-significant induction, respectively (Figure 6b). In line with the style elongation phenotype abundance and *Hsp17*.*6* induction, pollen viability tests further confirmed the heat stress tolerance of CLN1621 and Saladette. Under MCHS conditions, pollen viability rate was reduced to 3% of control in M82 and Moneymaker cultivars, while for CLN1621 and Saladette, pollen viability rates reduced to 43% and 22% respectively, compared with normal conditions (Figure 6c). Analyzing DPBA-stained pollen from these plants indicated that the tolerant cultivars accumulate about twice the amount of flavonols compared with the sensitive cultivars (Figure 6d).

**Figure 6.**
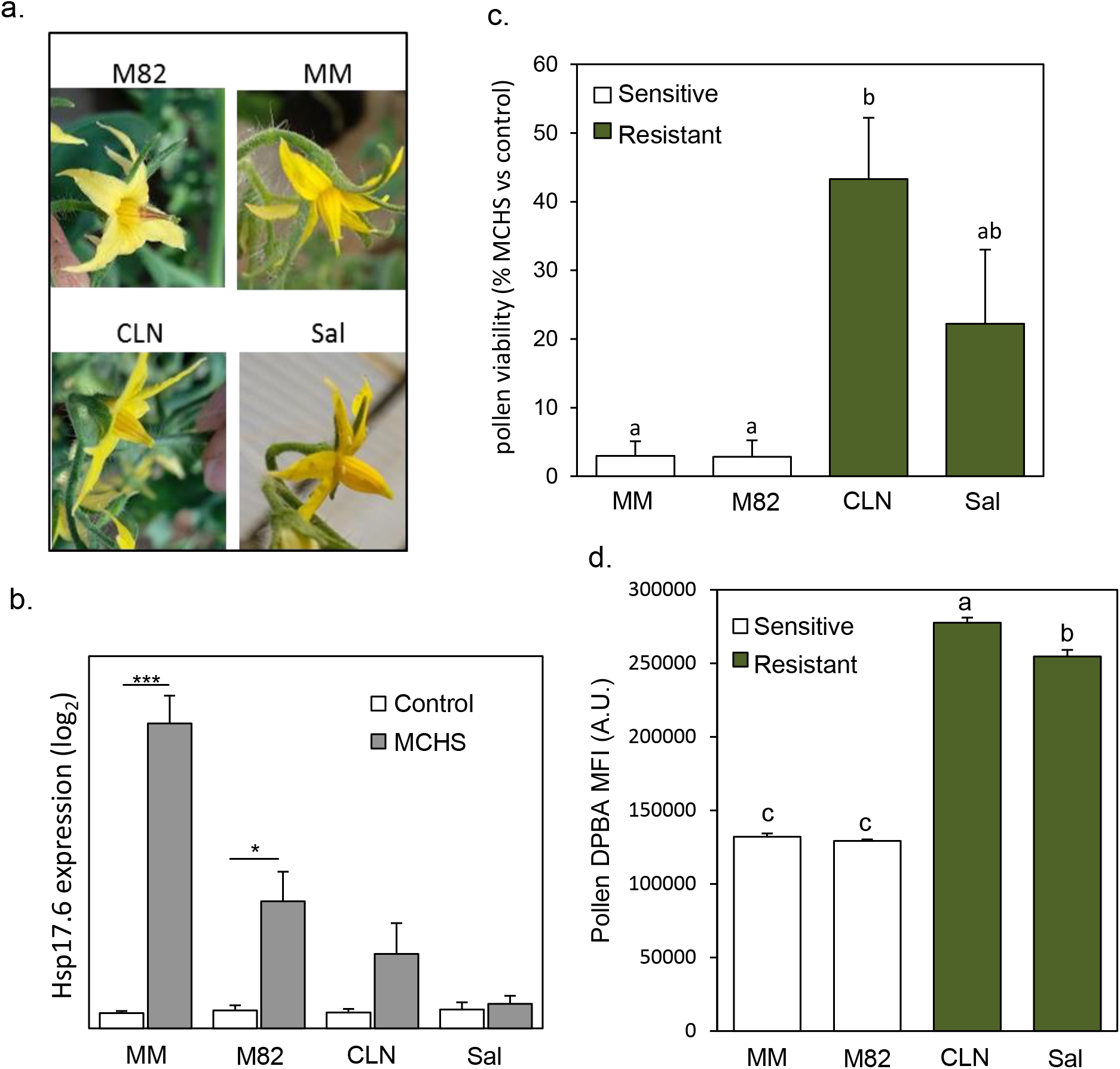
Flavonols levels are higher in pollen of heat stress tolerant tomato cultivars. (a) Typical flower appearance in response to heat stress in each of the four cultivars. (b) Expression level of the heat stress molecular marker gene *Hsp17*.*6* in leaves of the four genotypes, under control and MCHS conditions. (c) Pollen viability rates of the four genotypes under MCHS conditions, relative to control conditions. (d) Pollen flavonols in the four genotypes, measured by DPBA mean fluorescence signal intensity (MFI), in arbitrary units. *, p-value < 0.05. ***, p-value < 0.001. Different letters indicate statistically significant differences. MCHS, moderate chronic heat stress. MM, Moneymaker. CLN, CLN1621. Sal, Saladette.

## Discussion

Our findings indicate that the *hp2* tomato mutant is more tolerant than wild-type plants to conditions of moderate chronic heat stress (MCHS). We found that under these conditions, the *hp2* lines maintained a bigger fraction of viable pollen, with increased metabolic activity (Figure 2). This pollen fraction also kept its normal size under heat stress conditions, whereas the wild-type lines displayed a sharp decrease in the proportion of the functional, larger high-ROS pollen (Figure 3). Importantly, these findings were reproducible in three independent allelic lines of *hp2*, compared with two different isogeneic wild-type lines. By application of a short-term heat stress, we were able to identify better pollen germination in *hp2*^*dg*^ following the stress (Figure 4). These data show better thermotolerance of *hp2* specifically in pollen, compared with wild-type. The *hp2* lines maintained normal seeds production under MCHS, while the wild-type lines presented a sharp decrease in seeds set. We suggest that the improved pollen viability and normal seed set rate of *hp2* under MCHS conditions is, at least in part, due to the capability of *hp2* to accumulate higher levels of flavonols under heat stress conditions. Our finding that known thermotolerant tomato lines such as Saladette and CLN1621 have higher levels of pollen flavonols under normal conditions supports the assumption that flavonols are responsible for the improved thermotolerance of *hp2* pollen.

### Flavonols as heat stress ameliorating agents in pollen

Several studies pointed at the involvement of flavonols in the response to heat stress in plants. In tomato, flavonols content in leaves and pollen was increased about 2-fold in response to heat stress (Martinez et al. 2016; Paupière et al. 2017). Reduced levels of flavonols was functionally linked with a reduction in heat stress tolerance, as demonstrated by the heat stress sensitivity of the tomato *are* (anthocyanin reduced) mutant, which has lower levels of flavonols in pollen. In this mutant, pollen viability is strongly reduced when plants are grown under heat stress conditions (Muhlemann et al. 2018). The *hp2* mutant in tomato represents the opposite case, where flavonols content is elevated, offerings a unique opportunity for testing the effect of flavonols induction on pollen performance under heat stress conditions. Here we show for the first time that the *hp2* alleles increase pollen flavonols, adding to previous data showing increased flavonols in fruits of *hp2* (Bino et al. 2005). Interestingly, flavonols levels increased in both the wild-type and *hp2* pollen, although more dramatically in the latter (up to a 17.5-fold induction in *hp2*^*j*^), under MCHS conditions (Table S1), placing flavonols as a component of pollen heat stress response. These findings are in agreement with the function of flavonols as antioxidants and with previous findings showing their ability to protect against various environmental stresses such as heat, cold and drought (Muhlemann et al. 2018; Di Ferdinando et al. 2012; Nakabayashi et al. 2014; M. Zhao et al. 2019; Zandalinas et al. 2017). Thus, this antioxidative activity, may account, at least in part, to the improved ability of *hp2* to maintain pollen viability and seed setting under MCHS conditions (Figures 1 and 2).

Flavonols are highly abundant in the tapetum tissue of the anther and are transported as flavonol glycosides to the sporopollenin layer of the macrogametophyte during pollen development (Hsieh and Huang 2007; Fellenberg and Vogt 2015). Recent study in Arabidopsis identified FST1, a member of the nitrate/peptide transporter family that is expressed specifically in the tapetum and required for the accumulation and transport of pollen-specific flavonols to the pollen surface in Arabidopsis. In a mutant lacking this transporter, flavonols transport to the pollen is reduced, and pollen viability is hindered (Grunewald et al. 2020). Putative orthologous genes were identified in several plant species, including tomato. Our analysis of mature pollen flavonols do not distinguish flavonols originated from the sporopollenin surrounding the pollen grain, and flavonols that may come from the tapetum tissue breakdown during pollen maturation. Therefore, the induced level of flavonols we detected in pollen from *hp2* may be the result of either enhanced biosynthesis in the tapetum, enhanced transport to the sporopollenin, or both. The issues of spatial localization of flavonols biosynthesis and transport in *hp2* are yet to be investigated.

### Thermotolerant tomato cultivars

The cultivar Saladette was reported as heat-tolerant in 1977 (Rudich et al. 1977), and has been used as a heat-tolerant reference in numerous studies since. It was shown to cope better with high temperatures in terms of assimilate partitioning and metabolism, resulting in improved pollen viability and higher fruit set under heat stress conditions (Rudich et al. 1977; Dinar and Rudich 1985; Dinar et al. 1983; Abdul-Baki 1991; Firon et al. 2006).

CLN1621L was found to be heat stress tolerant in two independent screens for tolerant tomato cultivars (Comlekcioglu and Kemal Soylu 2010; Balyan et al. 2020). Under field conditions, when day/night temperatures were on average 37/27°C, CLN1621L produced a higher rate of seeded fruits and lower rate of aborted flowers compared with other cultivars tested (Comlekcioglu and Kemal Soylu 2010). Comparing different seasons, CLN1621L performed better in the hot season (day/night temperatures of 35/28°C) in terms of fruit set rate and number of pollen grains per flower (Sangu et al. 2015). Despite many publications characterizing the physiological response to heat stress in Saladette and CLN1621L, no data was available for pollen secondary metabolites. Our findings that both thermotolerant cultivars have about twice as much flavonols in their pollen compared with thermosensitive cultivars (Figure 6) provides perhaps the first mechanistic explanation, at least in part, for their increased reproductive success under high temperatures, and an independent support to our hypothesis. However, it will be interesting to conduct a controlled heat stress experiment similarly to the *hp2* system, in order to compare pollen flavonols levels between control and MCHS conditions.

### *hp2* mutants as a potential source for heat stress tolerance traits

The previously characterized phenotypes of *hp2* (e.g. increased accumulation of fruits carotenoids, flavonoids and vitamins) are caused by malfunction of the tomato *DEETIOLATED1* gene (*LEDET1*, Solyc01g056340). DET1 acts as a component of an E3 ubiquitin ligase complex targeting proteins for degradation by the 26S proteasome (Bernhardt et al. 2006; Zhang et al. 2008). Thus, DET1 is affecting the abundance of different genes in various biosynthetic pathways such as GLK2, BBX20 and MBD5, which act as upstream regulators of chloroplast biogenesis, flavonoid accumulation, and carotenoid biosynthesis in tomato (Li et al. 2016; Nguyen et al. 2014; Powell et al. 2012; Tang et al. 2016; Xiong et al. 2019). This dependency may explain the mode of action of the *hp2* mutant, suggesting that when DET1 function is impaired, the 26S proteasome-mediated degradation of positive regulators of plastids biogenesis and carotenoids biosynthesis is hindered, resulting in increased number of plastids and enhanced pigmentation. This work is the first evaluation, to our knowledge, of the effect of *hp2* secondary metabolites on pollen viability and performance. We show that the flavonols over-accumulation effect of *hp2* in fruit is reproduced in pollen (Figure 5). It is yet to be defined how DET1 function facilitates accumulation of flavonols in pollen under heat stress conditions. Based on current knowledge, we hypothesize that the function of the 26S proteasome in the tapetum is hindered when temperatures arise, leading to decreased degradation of positive regulators of flavonoids biosynthesis resulting in flavonols accumulation.

The marginal increase in pollen flavonols that we found under MCHS conditions in the wild-type lines Moneymaker and Manapal (Figure 5b) may suggest an additional pathway in which the response to heat stress could activate flavonols biosynthesis independently of the proteasome degradation machinery which involves DET1. To shed light on this issue, the function of the 26S proteasome under control and MCHS conditions should be tested.

Importantly, the beneficial effect of *hp2* in stress tolerance may extend beyond heat stress due to the fact that flavonols play an important role in protecting against oxidative stress which accompanies many other environmental stresses such as cold, drought and nutrient starvation (Nakabayashi et al. 2014; Song et al. 2020; M. Zhao et al. 2019; Zandalinas et al. 2017). Interestingly, a recent publication describes the tomato Micro-Tom *high-pigment 1*, a closely related mutant to *hp2*, as chilling stress tolerant in terms of photosynthetic activity, activities of antioxidant enzymes, and accumulation of protecting osmolytes (Shahzad et al. 2020). It would be of interest to test whether pollen characteristics such as viability, germination capacity and flavonol levels are playing a role also in *hp1* under chilling and heat stress conditions.

Finally, the enhanced accumulation of human-health promoting secondary metabolites drove the introgression of *hp2* mutations into commercial tomato cultivars, facilitating the production of lycopene-rich fresh market tomatoes as well as processing cultivars for cost efficient lycopene extraction for the food and cosmetics industries (Levin et al. 2006; Ilahy et al. 2018). The identification of *hp2* as a novel potential source for crop thermotolerance may increase its value as a biotechnology tool in plant breeding schemes. For this purpose, detailed characterization of yield traits such as fruit number, total weight and brix under control, MCHS and commercial growth conditions should be performed using the original *hp2* mutant lines (as in this study), as well as commercial cultivars that carry either of the *hp2* alleles in various backgrounds (Ilahy et al. 2018).

## Experimental procedures

### Seed set quantification

In order to determine seed set rate, ripe fruits from each plant were collected and sectioned. Fruits with no or little amount of seeds (<10 seeds) were considered as ‘unseeded fruits’, while fully seeded fruits were considered as ‘seeded’. Fruit pictures were taken using Cannon PowerShot SX520HS camera. For each plant, the ratio of seeded fruits out of total was calculated. For seed count, seeds were extracted using sulfuric acid. Seeds were extracted with the placenta cells and jelly tissues attached, incubated for 3 hours in a 2% sulfuric acid solution, then, transferred into net bags and rinsed with tap water. Clean seeds were left to air-dry, then spread over paper and imaged using a smartphone camera (exposure time 0.25 s, Mi6, Xiaomi, China). Seed number was quantified automatically by image processing in ImageJ (version 1.50i) using the “Count Kernels in Directory” macro as described in (Severini et al. 2011) with several modification and adjustments (circularity 0.1-1.00, size>=150).

### Pollen germination assays

To isolate pollen for *in vitro* germination, individual anther cones were detached from open flowers, cut in transverse and placed in 1.2ml of pollen germination medium (2 mM boric acid, 2 mM calcium nitrate, 2 mM magnesium sulfate and 1 mM potassium nitrate, suplemented with 10% w/v sucrose before use). Cut anther cones in solution were gently crushed and vortexed to release pollen into the medium, and then split into two 1.5 ml Eppendorf tubes each of 0.5 ml pollen suspension. One tube was then incubated at 34°C for 30 min (for *in vitro* heat stress), while the other tube was incubated at 22°C for 30 min (as control). Pollen (0.1 ml) from each condition was then transferred to a 96-well plate, covered and incubated at 22°C for 1.5 hours. Pollen were observed using an Olympus IX2 inverted microscope and images were captured at X4 magnification using an Olympus XM10 camera. Scoring of germination was performed manually using the ImageJ (version 1.50i) Cell Counter plug-in (Schneider et al. 2012).

### Isolation of pollen for flow cytometry

To isolate pollen, 3-4 anther cones were detached from open flowers, cut in transverse and placed in 5 ml of 10% sucrose pollen germination medium in a 15-ml centrifuge tube. Pollen was released from anther cones by gently crushing against the wall of the tube followed by vortexing for 30 seconds. Suspensions of pollen were subsequently filtrated through folded miracloth (Merck Millipore, Darmstadt, Germany) to remove tissue debris.

### Staining pollen with H_2_DCFDA or DPBA for flow cytometry

H_2_DCFDA and DPBA were dissolved in DMSO. Pollen was allowed to hydrate in germination buffer for 15 min before H_2_DCFDA (Cayman chemical) or DPBA (2-aminoethyl diphenylborinate; Sigma) was added to a final concentration of 5 µM and 0.025 %w/v, respectively. Pollen were incubated in stains for 30 min prior to flow cytometry analysis. A sample of unstained pollen was retained to calibrate the DCF and DPBA fluorescence threshold. Mode of action of H_2_DCFDA: Upon hydrolysis by cytoplasmic esterases, the DA moiety of H_2_DCFDA is cleaved-off, freeing the non-fluorescent H_2_DCF to become oxidized by ROS, forming green fluorescent dichlorofluorescein (DCF). Thus, only metabolically active cells are DCF-stained, and the mean fluorescent intensity (MFI) signal correlates with the rate of metabolic activity and level of ROS in the cell. Therefore, DCF staining is used to measure cell viability (Rutley and Miller 2020; Luria et al. 2019).

### Flow cytometry analyses of stained pollen grains

Following staining, samples were analyzed in a BD LSRFortessa™ and the FITC laser-filter set (488 nm solid state laser, BP filter 530/30) compatible with both DCF and DPBA. Gating of singlet pollen grains was performed by first gating pollen from non-pollen events using Forward Scatter Area (FSC-A) and Side Scatter Area (SSC-A) parameters, and then applying a gate for double discrimination using Forward Scatter Height (FSC-H) and Forward Scatter Width (FSC-W). In the FITC-A channel, true fluorescence was gated from auto-fluorescence using the unstained sample as the auto-fluorescence threshold. Typically, > 3000 singlet pollen grains were analyzed per sample. Data analyses were performed using FlowJo software.

### ImageStream®X Mk II imaging flow cytometry

Reference images of DCF and DPBA-stained pollen from Moneymaker and *hp2* were acquired using Amnis® ImageStream®X Mk II (MilliporeSigma) imaging flow cytometer and INSPIRE (MilliporeSigma) software was used for data collection. Samples were acquired at 40X magnification and medium spend/medium resolution. The bright field images were acquired in Channel 1 and DCF or DPBA signal in Channel 2 (488 nm laser, BP filter 528/65). Image analyses were performed using the IDEAS® software.

### RNA isolation and quantitative-PCR analysis

Anther cones were detached from open flowers and flash frozen in liquid nitrogen. Total RNA was isolated from pairs of anther cones using RiboEx™ (GeneAll; Seoul, South Korea) according to the manufacturer’s protocol. RNA was treated with DNaseI (Thermo Fischer Scientific, California, USA) prior to cDNA synthesis. cDNA was synthesized from 0.5 µg of DNaseI-treated total RNA using qPCRBIO cDNA Synthesis Kit (PCR Biosystems, London, United Kingdom) following the manufacturer’s instructions. Relative expression of HSP17.6 (NM_001246984.3) and the reference genes ubiquitin (Solyc07g064130) for Moneymaker, *hp2* and *hp2*^*j*^ or EF1α (Solyc06g009970) for *hp2*^*dg*^ and Manapal were determined using quantitative real-time PCR (qRT-PCR) on an Applied Biosystems StepOnePlus Real-Time PCR System (Thermo Fischer Scientific, California, USA). Each 10 µl qRT-PCR reaction consisted of qPCRBIO SyGreen Blue Mix Hi-ROX (PCR Biosystems, London, United Kingdom), gene-specific primers and 1:5 diluted cDNA template. Thermocycling conditions were performed according to manufacturer instructions. Data were analyzed using Applied Biosystems *StepOne*™ software v 2.3 (Thermo Fischer Scientific, California, USA). Primer sequences are listed in Table S3.

## Acknowledgments

This research was supported by institutional grants to MLL and grants to GM and JFH from BARD IS-4652-13, GM from BSF-2016605, JFH from NSF IOS 1656774

## Abbreviations

MCHS: Moderate Chronic Heat Stress
WT: Wild-type
hp: high-pigment
MM: Moneymaker
MP: Manapal
MT: MicroTom
CHS: chalcone synthase
CHI: chalcone isomerase
ROS: reactive oxygen species
H_2_DCFDA: 2’,7’-dichlorodihydrofluorescein diacetate
DCF: dichlorofluorescein
DPBA: diphenylboric acid-2-aminoethyl ester
MFI: Mean Fluorescent Intensity

