## Supporting information for "Enhanced reproductive thermotolerance is associated with increased accumulation of flavonols in pollen of the tomato *high-pigment* 2 mutant"

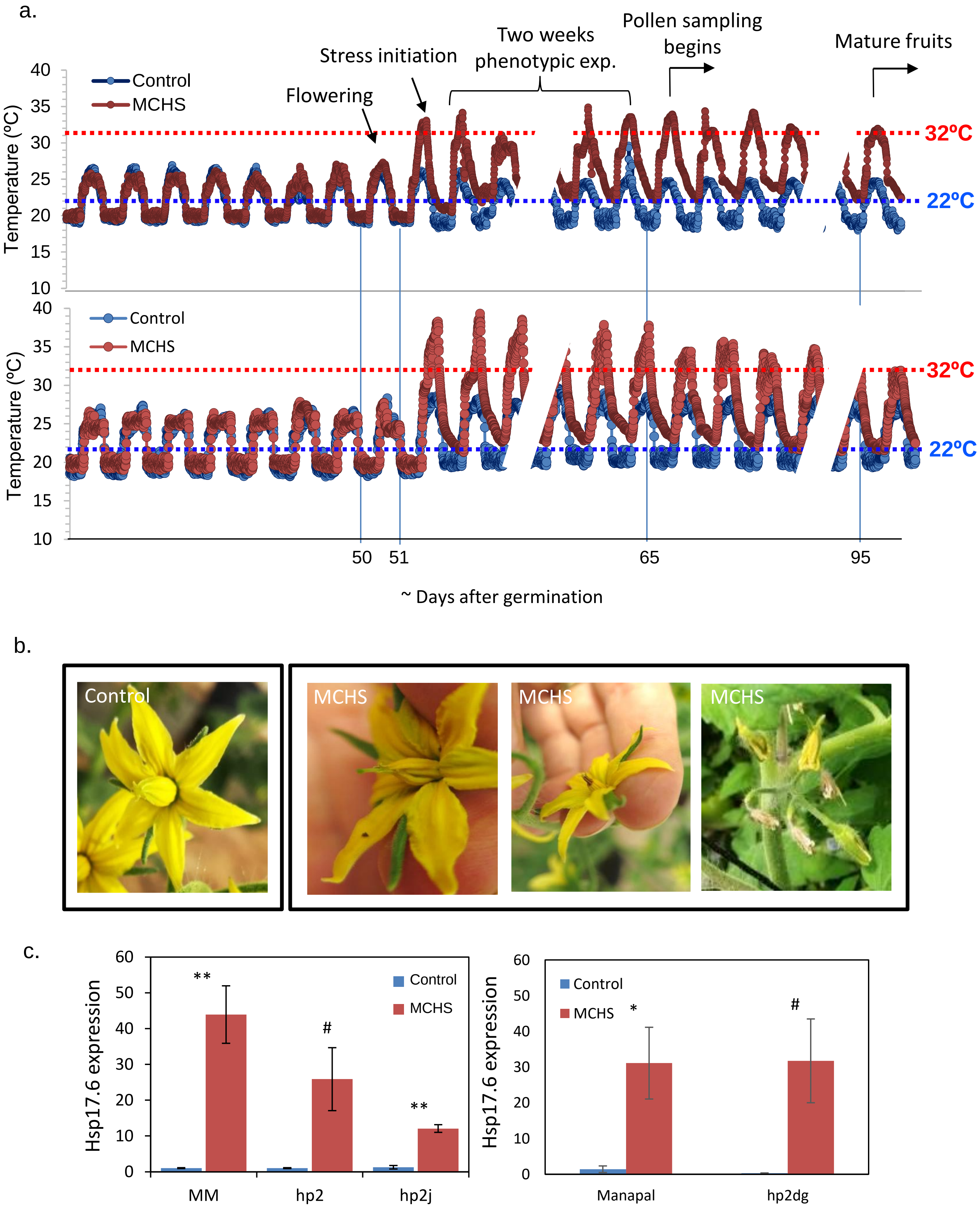


**Figure S1. Experimental conditions and flowers phenotypes.** (a) Recorded temperatures in degrees Celsius every 10 minutes in the control (blue) and MCHS (brown) greenhouses several days before and after stress initiation. Upper panel: first experiment, April-July 2019. Lower panel: second experiment, August-November 2019. (b) Flower appearance in the control conditions greenhouse (left) versus flower phenotypes under MCHS conditions (right). (c) Expression level of the heat stress marker gene *Hsp17.6* under control and MCHS conditions. *, p-value <0.05. **, p-value <0.01. ***, p-value <0.001. # p-value < 0.055.

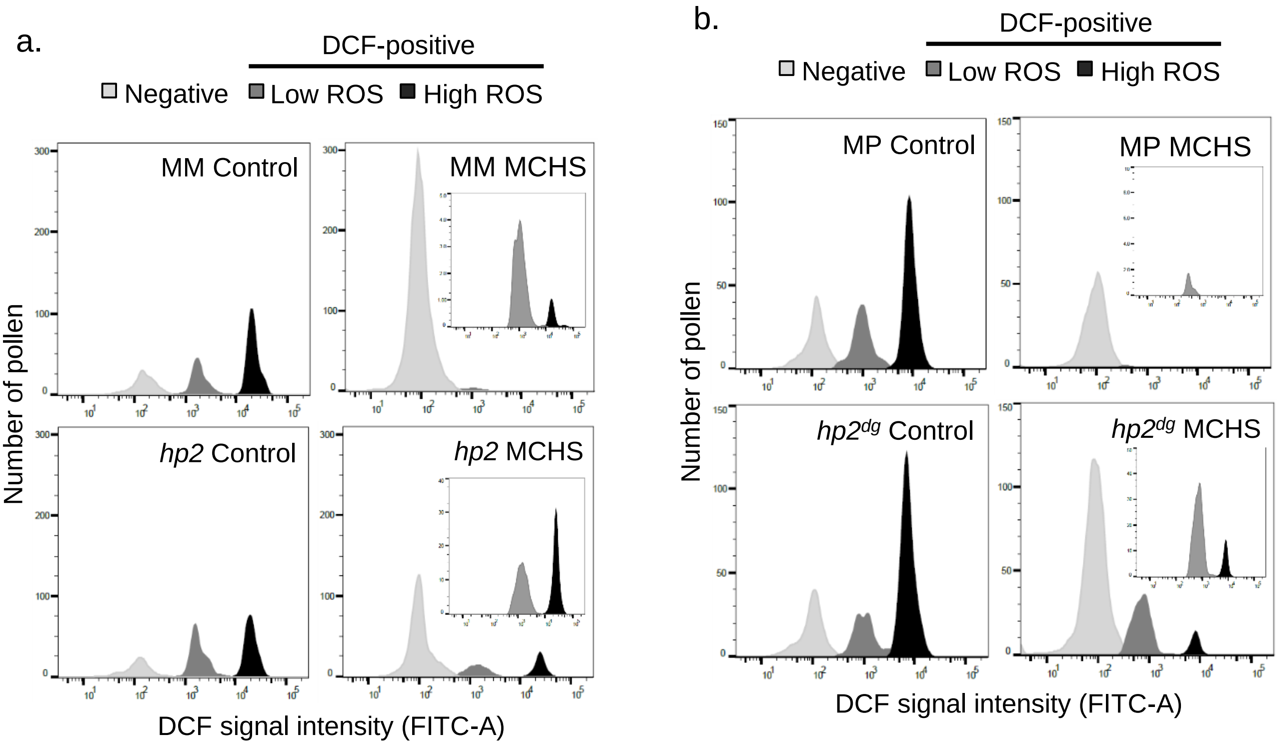


**Figure S2. Pollen DCF signal intensity distribution plots for *hp2* and *hp2*$^{dg}$ genotypes.** Inner frames zoom into the DCF-positive fraction, which is smaller under MCHS conditions in all genotypes. MCHS, moderate chronic heat stress. MM, Moneymaker. MP, Manapal.

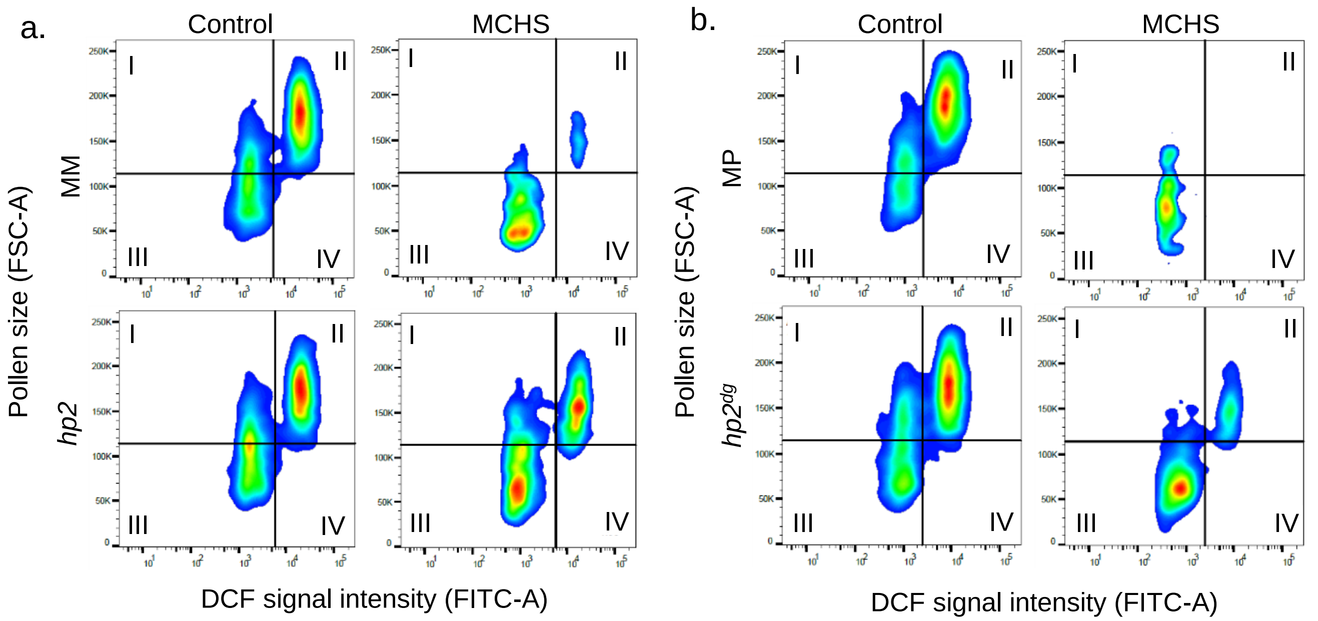


**Figure S3. Pollen grain size (FSC-A channel, Y-axis) versus pollen ROS level (FITC-A signal, X axis) for positive DCF pollen in *hp2* and *hp2*$^{dg}$ genotypes.** The larger, high-ROS pollen faction is represented in quadrant II**.** MCHS, moderate chronic heat stress. MM, Moneymaker. MP, Manapal.

a.

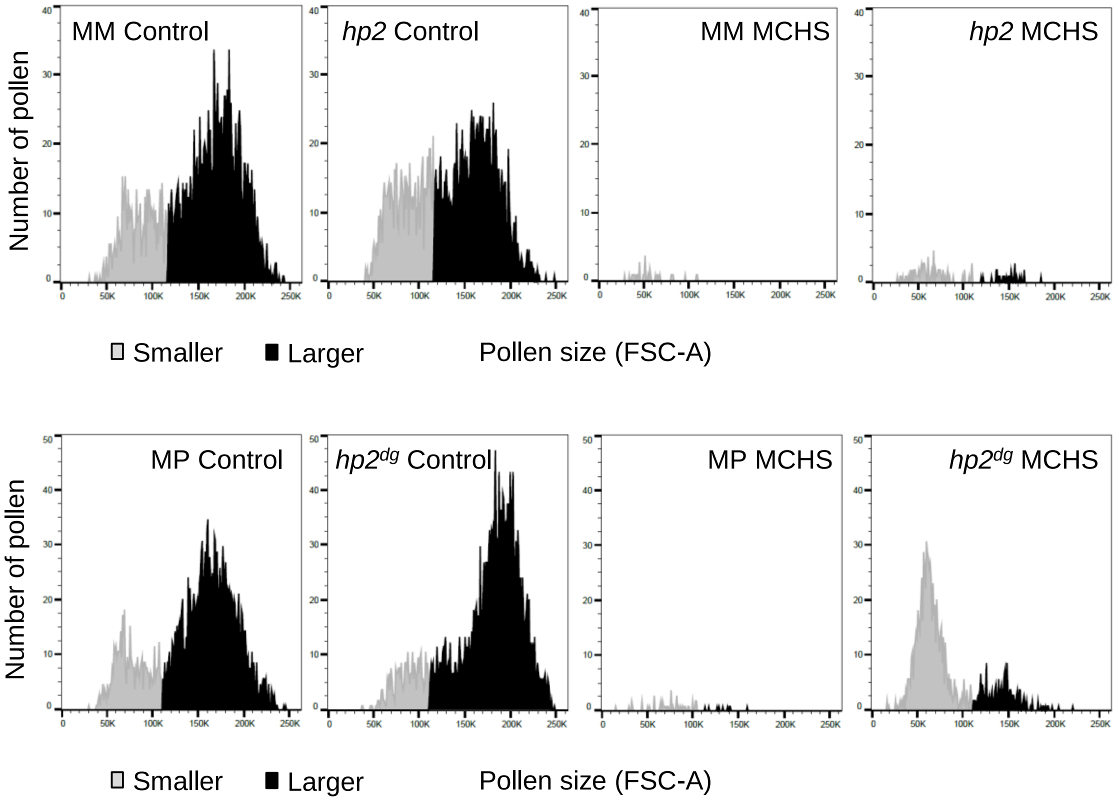


**Figure S4. DCF-positive pollen size distribution for *hp2* and *hp2*$^{dg}$ genotypes.** MCHS, moderate chronic heat stress. MM, Moneymaker. MP, Manapal.

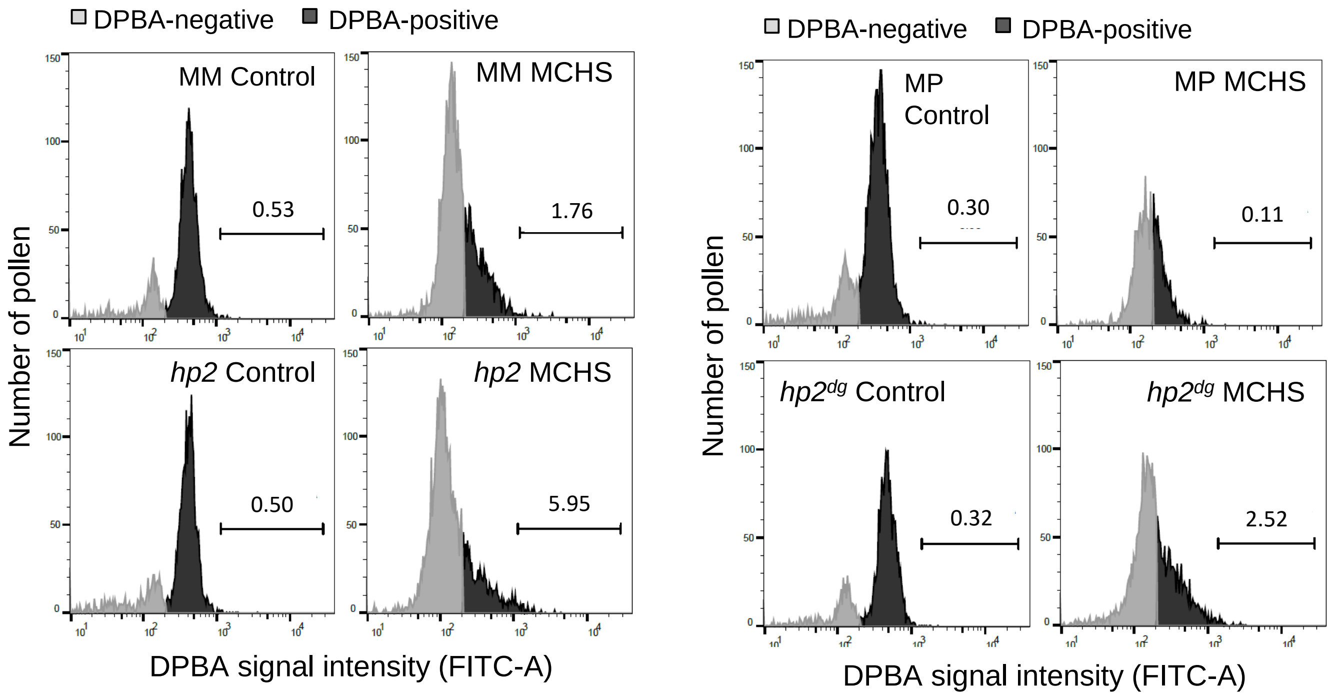


**Figure S5. DPBA-stained pollen signal intensity distribution for *hp2* and *hp2[dg]* genotypes.** MCHS, moderate chronic heat stress. MM, Moneymaker. MP, Manapal.

### Table S1: Raw DCF and DPBA values and fold change

| | | | Flavonols assay - DPBA signal | | | | | | Viability assay - DCF signal | | | |
|---|---|---|---|---|---|---|---|---|---|---|---|---|
| Exp. # | line | condition | MFI | mutant vs. wild type | MCHS vs. control | % enhanced | mutant vs. wild type | MCHS vs. control | pollen population | % | mutant vs. wild type | MCHS vs. control |
| 1 | Moneymaker (MM) | control | 425.4 | | | 0.5 | | | no ROS | 25.2 | | 3.79 |
| | | | | | | | | | low ROS | 21.8 | | 1.22 |
| | | | | | | | | | high ROS | 53.1 | | 1.00 |
| | | MCHS | 438.8 | | 1.03 | 3.3 | | 6.60 | no ROS | 95.6 | | |
| | | | | | | | | | low ROS | 26.6 | | |
| | | | | | | | | | high ROS | 53.1 | | |
| | *hp2* | control | 417.0 | 0.98 | | 0.5 | 1.00 | | no ROS | 20.3 | 0.81 | 3.44 |
| | | | | | | | | | low ROS | 26.6 | 1.22 | 0.44 |
| | | | | | | | | | high ROS | 53.1 | 1.00 | 0.35 |
| | | MCHS | 518.6 | 1.18 | 1.24 | 5.8 | 1.76 | 11.60 | no ROS | 69.9 | 0.73 | |
| | | | | | | | | | low ROS | 11.7 | 0.44 | |
| | | | | | | | | | high ROS | 18.4 | 0.35 | |
| 2 | Moneymaker (MM) | control | 402.0 | | | 0.3 | | | no ROS | 27.7 | | 3.49 |
| | | | | | | | | | low ROS | 29.5 | | 0.11 |
| | | | | | | | | | high ROS | 42.8 | | 0.00 |
| | | MCHS | 440.7 | | 1.10 | 3.8 | | 12.67 | no ROS | 96.8 | | |
| | | | | | | | | | low ROS | 3.1 | | |
| | | | | | | | | | high ROS | 0.1 | | |
| | $hp2^{j}$ | control | 504.3 | 1.25 | | 2.0 | 6.67 | | no ROS | 24.4 | 0.88 | 3.53 |
| | | | | | | | | | low ROS | 34.5 | 1.17 | 0.32 |
| | | | | | | | | | high ROS | 41.0 | 0.96 | 0.07 |
| | | MCHS | 800.0 | 1.82 | 1.59 | 35.0 | 9.21 | 17.50 | no ROS | 86.2 | 0.89 | |
| | | | | | | | | | low ROS | 11.1 | 3.58 | |
| | | | | | | | | | high ROS | 2.7 | 27.00 | |
| 3 | Manapal (MP) | control | 487.2 | | | 0.4 | | | no ROS | 28.2 | | 3.49 |
| | | | | | | | | | low ROS | 27.3 | | 0.06 |
| | | | | | | | | | high ROS | 44.5 | | 0.00 |
| | | MCHS | 522.4 | | 1.07 | 0.7 | | 1.75 | no ROS | 98.4 | | |
| | | | | | | | | | low ROS | 1.6 | | |
| | | | | | | | | | high ROS | 0.0 | | |
| | $hp2^{dg}$ | control | 556.8 | 1.14 | | 0.3 | 0.75 | | no ROS | 28.9 | 1.02 | 2.79 |
| | | | | | | | | | low ROS | 19.2 | 0.70 | 0.80 |
| | | | | | | | | | high ROS | 51.9 | 1.17 | 0.08 |
| | | MCHS | 624.0 | 1.19 | 1.12 | 1.0 | 1.43 | 3.33 | no ROS | 80.5 | 0.82 | |
| | | | | | | | | | low ROS | 15.3 | 9.56 | |
| | | | | | | | | | high ROS | 4.2 | na | |

Table S2: Pearson’s correlation coefficient between FITC-A and FSC-A signal intensities for DCF-positive pollen.

| Line | Condition | N | Corr coeff |
|---|---|---|---|
| MM | Control | 6 | 0.77 |
| | MCHS | 6 | 0.67 |
| *hp2* | Control | 6 | 0.78 |
| | MCHS | 6 | 0.78 |

| | | | |
|---|---|---|---|
| MM | Control | 3 | 0.76 |
| | MCHS | 3 | 0.52 |
| *hp2$^{j}$* | Control | 3 | 0.76 |
| | MCHS | 3 | 0.81 |

| | | | |
|---|---|---|---|
| MP | Control | 3 | 0.70 |
| | MCHS | 3 | 0.75 |
| MP *hp2$^{dg}$* | Control | 3 | 0.78 |
| | MCHS | 3 | 0.77 |

### Table S3: Primer sequences for qRT-PCR

| **Name** | **Gene** | **Forward** | **Reverse** |
|---|---|---|---|
| HSP17.C | Solyc08g062450 | GGAAGAGGGAAGAAGAGAAAGAA | ACCACAAACCATCAAAACAGAGT |
| UBI (reference) | Solyc07g064130 | GGACGGACGTACTCTAGCTGA T | AGCTTTCGACCTCAAGGGTA |
| eEF1 | Solyc06g009970 | AGTCAACTACCACTGGTCAC | GTGCAGTAGTACTTAGTGGTC |
